# Nano-analytical Characterization of Breast Tissue Biopsies Reveals an Association between Spherical Nano and Micro Particles and Invasive Malignant Breast Tumours

**DOI:** 10.1101/2020.04.29.067660

**Authors:** Elena Tsolaki, William Doran, Jan Overbeck, Luca Magnani, Alessandro Olivo, Inge K. Herrmann, Sergio Bertazzo

## Abstract

The presence of calcifications on mammograms is a widely used diagnostic marker for breast cancer. While the clinicopathological analysis of breast tissue is well-developed, an in-depth characterisation of the properties of calcifications at the micro and nano scale remained elusive. In this work, we use nano-analytical methods to investigate the calcification present in healthy breast tissue, as well as in benign and malignant breast tumour tissue biopsies. While polycrystalline apatite with lower crystallinity can be found in all breast tissue samples, we show that magnesium-containing nano and micro spherical particles are found only in malignant invasive breast cancers. These particles concentration increases with cancer progression. The discovery of these spherical particles provides new insights into the characteristics of breast calcification associated with malignant tumours and opens new research pathways to a better understanding of breast tumours and their microenvironment in general.

## Introduction

Calcification formed from mineral deposits with sizes in the hundreds of micrometres (known as microcalcifications) found in breast tissue are a crucial component in the diagnosis of underlying breast carcinomas^1^ and are commonly used in the differentiation between malignant and benign tumours^1-3^. Multiple similar calcification clusters, appearing as bright white spots on mammograms, spreading over one or both breasts are considered to be concomitant of benign tumours, whereas variability in the appearance of clusters suggests a malignant cause^1^.

This connection between breast calcification and cancer has led to many research initiatives aiming to identify specific differences between benign and malignant calcification^4-6^. Due to the resolution limit of clinical mammography, however, it is not possible to gain in-depth information on the mineral forming these calcifications. Similarly, the low resolution of both mammography and light microscopy used for the histopathological assessment of breast biopsies does not allow characterization at the nano scale, nor does it provide any information on chemical or physical properties of these calcifications.

In the past, characterisation methods typically used in materials sciences, such as X-ray, electron diffraction, electron microscopy, and microprobe analysis, have been applied to the calcification in breast cancer and suggested the presence of the mineral hydroxyapatite^4,7-10^. Hydroxyapatite has also been recorded widely in studies using spectroscopic and diffraction methods, in both benign and malignant cases^4,11^. The crystallinity of the hydroxyapatite mineral observed was found to be correlated with tumour invasiveness, with a higher crystallinity associated with malignant invasive tumours^12^. Recent studies, using Raman and energy dispersive x-ray spectroscopy, have also suggested the presence of magnesium-substituted apatite and whitlockite in malignant cases; however, no definite association has been reported^7,8,10^.

Other than their diagnostic potential, the physicochemical properties of the minerals present in the calcifications in breast cancer are of growing interest due to their potential for providing insights on the prognosis of the disease^13,14^. It has been reported that breast cancer patients with small calcifications have a lower survival rate than patients who present more extensive calcifications^15^, and that the presence of calcifications in milk ducts increases the risk of cancer recurrence^14^. Studies have also reported that the presence of hydroxyapatite in breast cancer cell cultures can enhance the processes of mitosis^16^ and the migration of tumour cells^17^. Furthermore, tumorigenic mammary cells can reportedly produce hydroxyapatite, in contrast to non-tumorigenic cells, which do not calcify^18^.

Despite the advances made in the field, the exact relationships between the physicochemical characteristics of the minerals present in different types of cancerous calcification and their association to cancer progression or prognosis are not fully established. A more comprehensive understanding of these may, therefore, shed light on the general nature of the calcification in tumours, and allow a correlation between calcification and tumour key characteristics.

Here we analyse histological slides obtained from breast tissue biopsies of eighty one patients with benign and malignant tumours of all grades, using nano-analytical methods such as Focused Ion Bean (FIB), Scanning and Transmission Electron microscopy (SEM and TEM), Selected Area Electron Diffraction (SAED), Energy-dispersive X-ray spectroscopy (EDS), and Raman spectroscopy. We show the distinct mineral characteristics of breast tissue calcifications present in benign and malignant tumours, and reveal that the presence of magnesium-rich spherical particles is directly associated with invasive malignant breast tumours.

## Results and Discussion

Density Dependent Colour-SEM (DDC-SEM) images show that calcification was present in two distinct morphologies. Large minerals (Fig. 1a, b and c) with no specific shape (dimensions between 10 and 300 µm) were observed in all tissue types (healthy, benign and malignant). Nano and micro spherical particles (Fig. 1d, e and f) with dimensions from 46 nm to 2.15 µm (size distribution in Supplementary Fig. S1) were only observed in invasive malignant cases.

**Figure 1:**
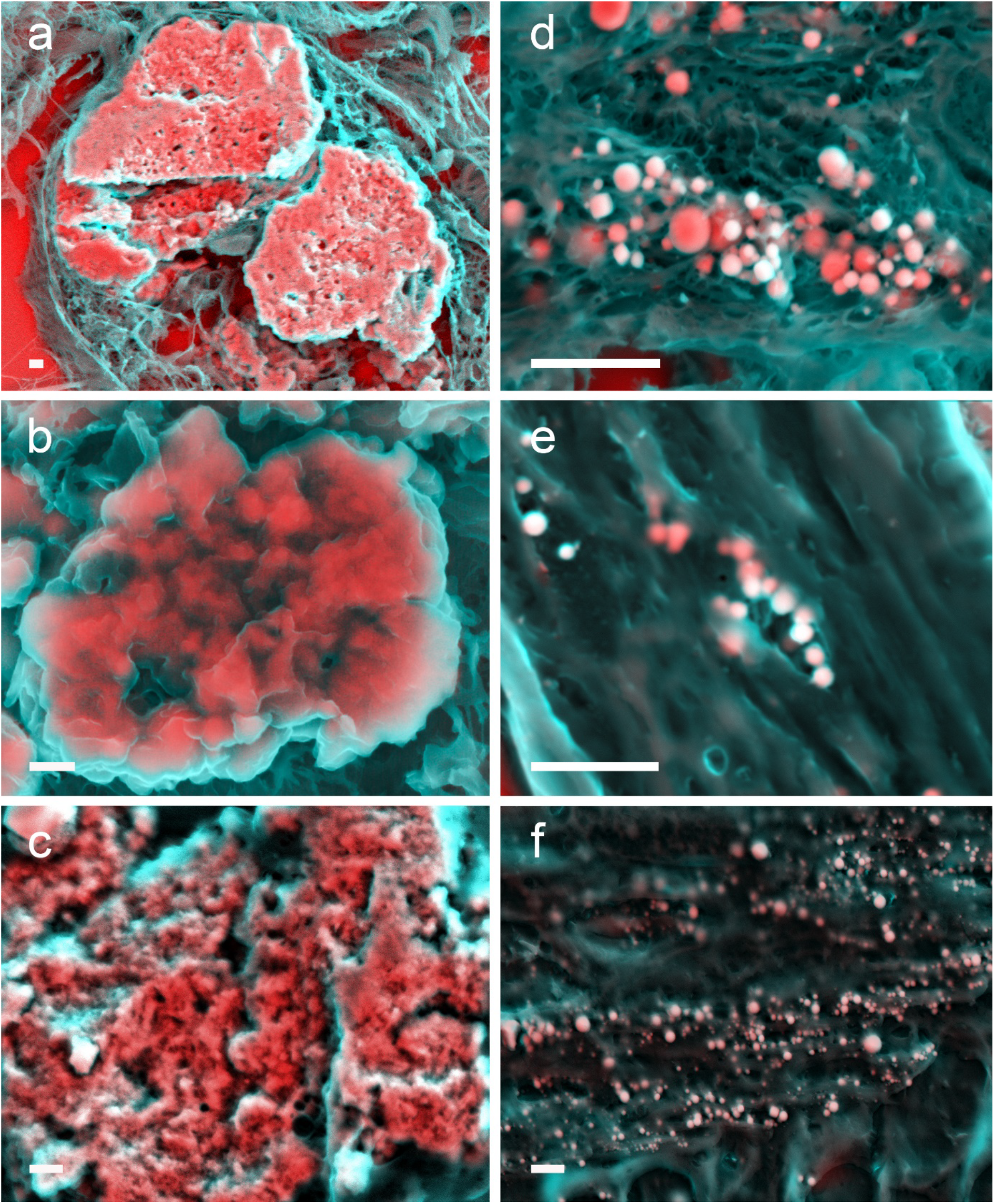
Density dependent colour scanning electric micrographs (DDC-SEM) of calcification observed in breast tissue. (**a**) DDC-SEM micrograph of large minerals present in a benign borderline phyllodes tumour. (**b**) DDC-SEM micrograph of large minerals present in a benign sclerosing papillary case. (**c**) DDC-SEM micrograph of large minerals present in a healthy tissue sample. (**d**) DDC-SEM micrograph of spherical particles in an invasive lobular carcinoma biopsy. (**e**) DDC-SEM micrograph of spherical particles in an invasive ductal carcinoma biopsy. (**f**) DDC-SEM micrograph of spherical particles in an invasive cribriform carcinoma biopsy. Red and pink indicate inorganic material and turquoise (blue/green) indicates organic material. Scale bar = 2 µm.

EDS analysis shows that all large minerals were composed of calcium and phosphorus, without a detectable amount of magnesium (Fig. 2a and b). The spherical particles, instead, were composed of calcium, phosphorus and magnesium (Fig. 3a, b and c).

**Figure 2:**
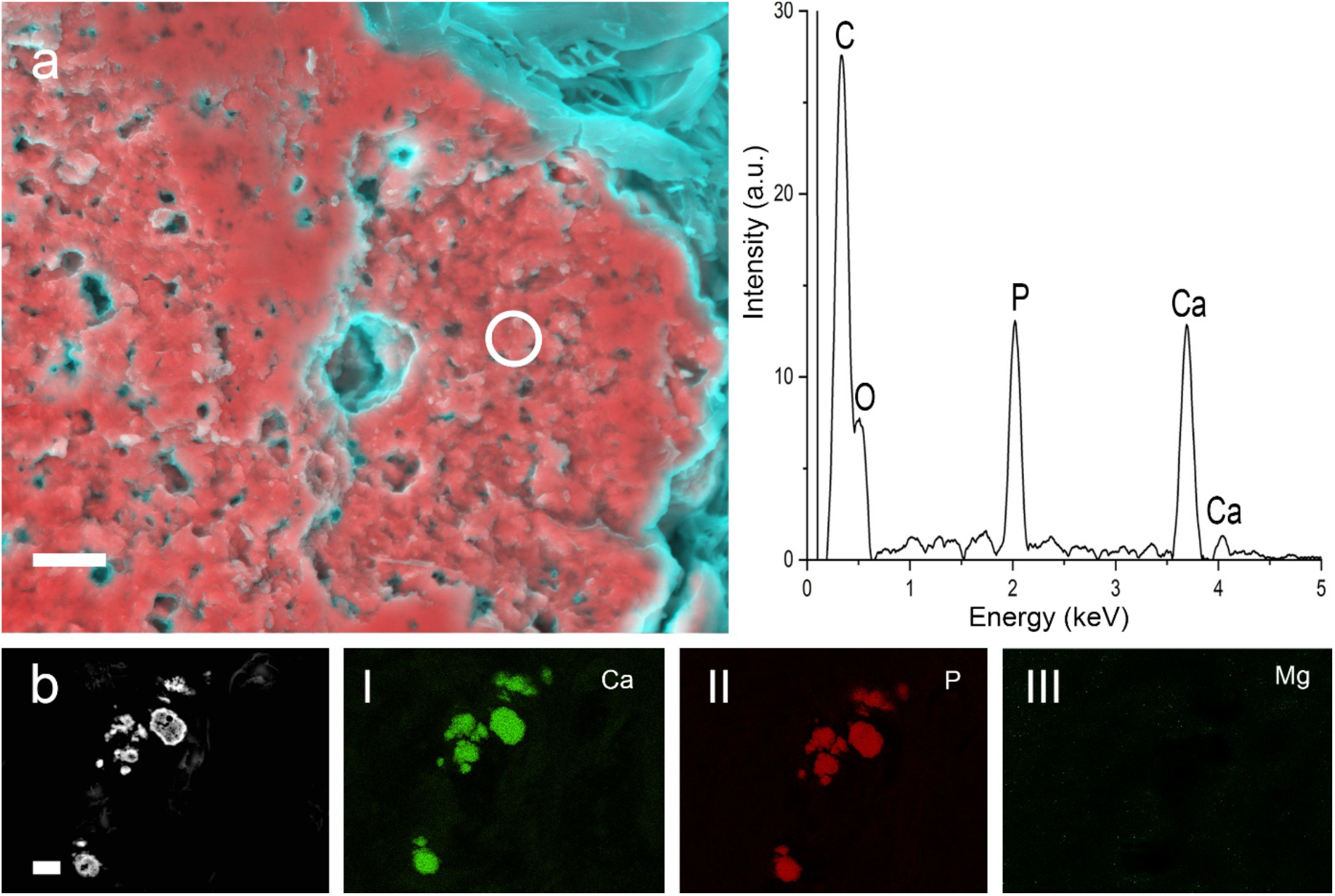
DDC-SEM micrographs and elemental mapping EDS spectra of large minerals found in benign breast tissue tumours. (**a**) DDC-SEM micrograph of large mineral present in a benign borderline phyllodes with EDS spectrum of the area marked with a circle. Red and pink indicate inorganic material and turquoise (blue/green) indicates organic material. Scale bar = 2 µm. (**b**) Scanning electron backscattering micrograph of large minerals in a benign complex sclerosing papillary lesion, with EDS elemental mapping of (**I**) calcium, (**II**) phosphorus and (**III**) magnesium. Scale bar = 10 µm.

**Figure 3:**
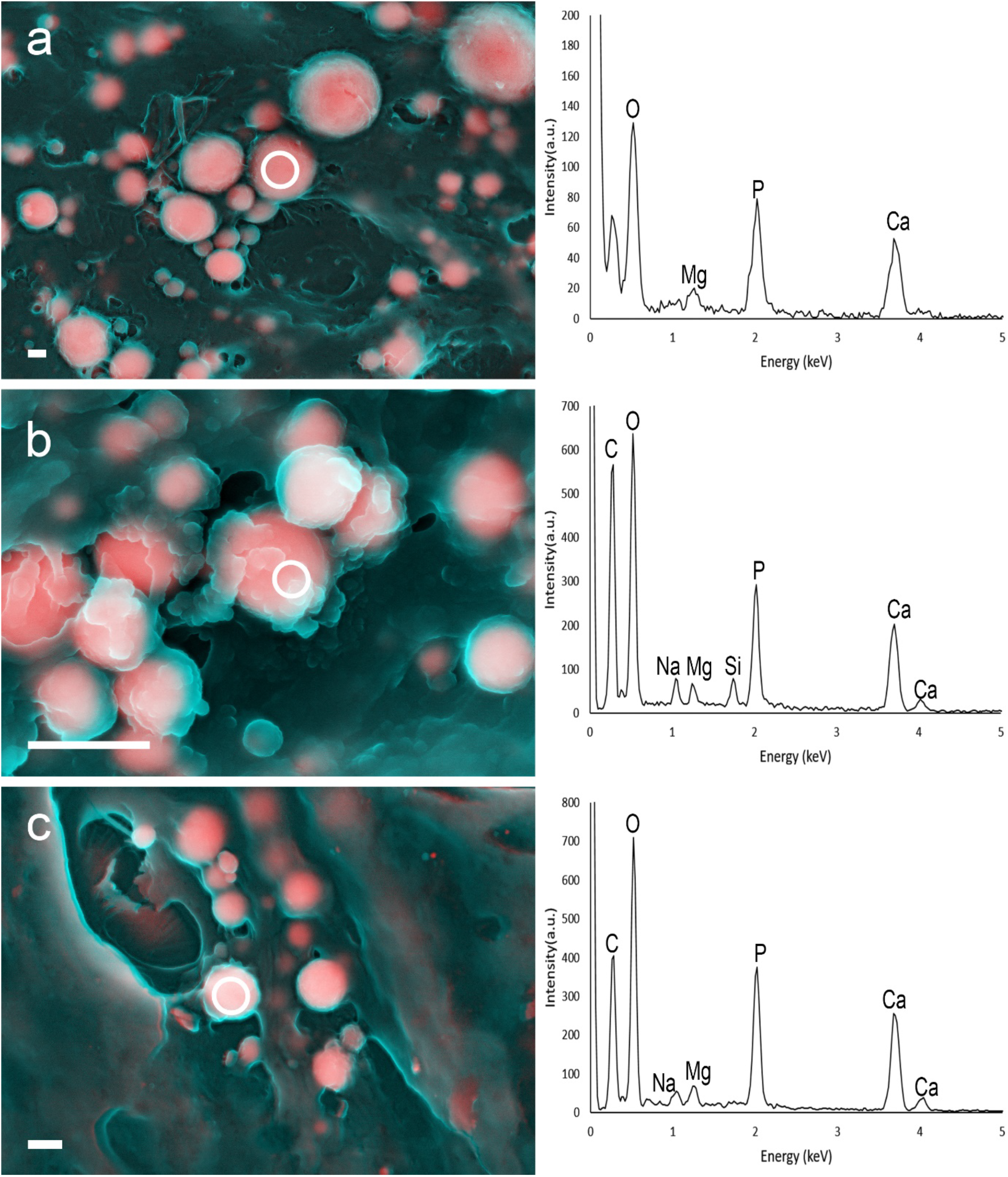
DDC-SEM micrographs and EDS spectra of spherical particles found in invasive malignant breast tumours. (**a**) DDC-SEM micrograph of spherical particles present in an invasive cribriform carcinoma, with EDS spectrum of the area marked by circle. (**b**) DDC-SEM micrograph of spherical particles present in an invasive lobular carcinoma biopsy, and EDS spectrum of the area marked with a circle. (**c**) DDC-SEM micrograph of spherical particles present in an invasive ductal carcinoma, and EDS spectrum of the area marked with a circle. Scale bar = 500 nm. Red and pink indicate inorganic material and turquoise (blue/green) indicates organic material.

To investigate their internal structure and crystallinity, the minerals were sectioned by FIB and imaged by TEM. For the large minerals, TEM micrographs revealed an unorganised internal structure (Fig. 4a). The SAED patterns shows typical rings formed from a polycrystalline mineral (Fig. 4b) composed of apatite (SAED indexing for apatite in Supplementary Fig. S2)^19^.

**Figure 4:**
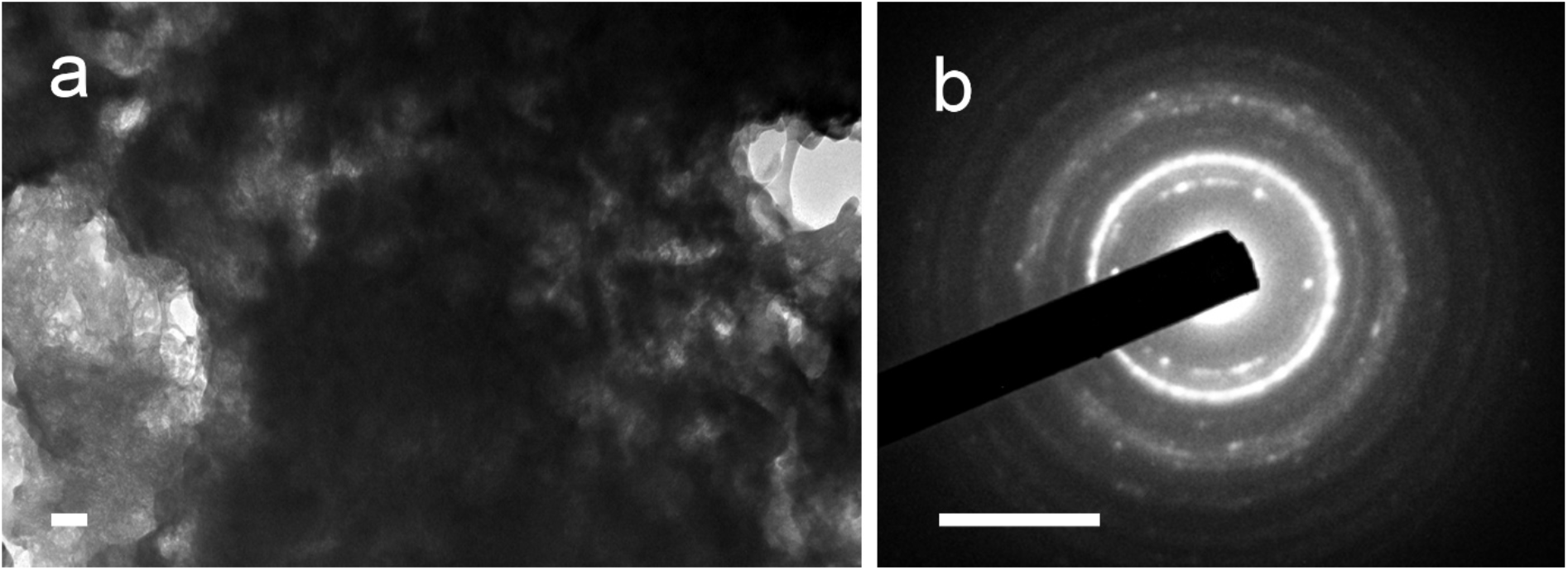
TEM micrograph of large mineral from benign phyllodes tumour sectioned by FIB, and representative SAED pattern. (**a**) TEM micrograph of large mineral. Scale bar = 100 nm. (**b**) SAED pattern of the region in **a**. Scale bar = 5 nm^-1^.

Further TEM micrographs of FIB-sectioned spherical particles (Fig. 5) show that their internal structure is heterogeneous, with some showing less dense cores (Fig. 5a, b and d). The respective SAED patterns, on the other hand, present with clear spots rather than rings (Fig. 5g, h, i). Indexing of the diffraction patterns suggests that the particles could be composed of whitlockite (SAED indexing for whitlockite in Supplementary Fig. S3)^20^.

**Figure 5:**
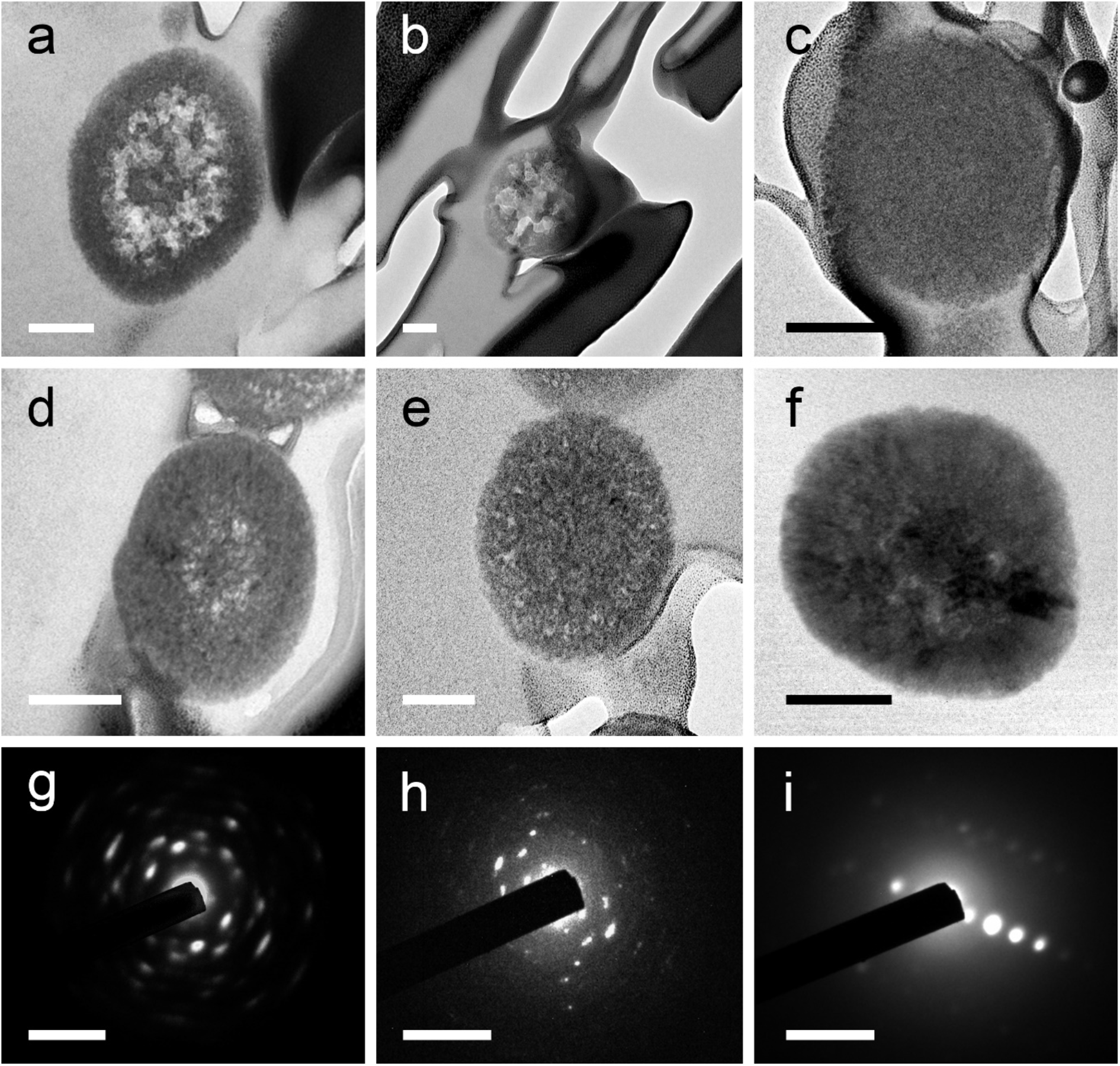
TEM micrographs of FIB-prepared sections of spherical particles and SAED patterns. (**a**) TEM micrograph of a spherical particle from invasive ductal carcinoma biopsy. (**b**) TEM micrograph of spherical particle from invasive ductal carcinoma biopsy. (**c**) TEM micrograph of spherical particle from invasive cribriform carcinoma biopsy (**d**) TEM micrograph of spherical particles from invasive lobular carcinoma biopsy. (**e**) TEM micrograph of spherical particles from invasive lobular carcinoma biopsy. (**f**) TEM micrograph of a spherical particle from invasive lobular carcinoma biopsy. Scale bars = 100 nm. (**g**) SAED pattern of the particles shown in **d**. (**h**) SAED pattern of the particles shown in **e**. (**i**) SAED pattern of the particles shown in **f**. Scale bars = 5 nm^-1^.

Finally, the chemical identity of both mineral types (large minerals and spherical particles) was further supported via Raman spectroscopy measurements. A characteristic peak at 960 cm^-1^ for apatite was observed for the large minerals (Fig. 6), and a peak at 970 cm^-1^, typical of whitlockite (Fig. 6), was observed for the spherical particles^21-23^.

**Figure 6:**
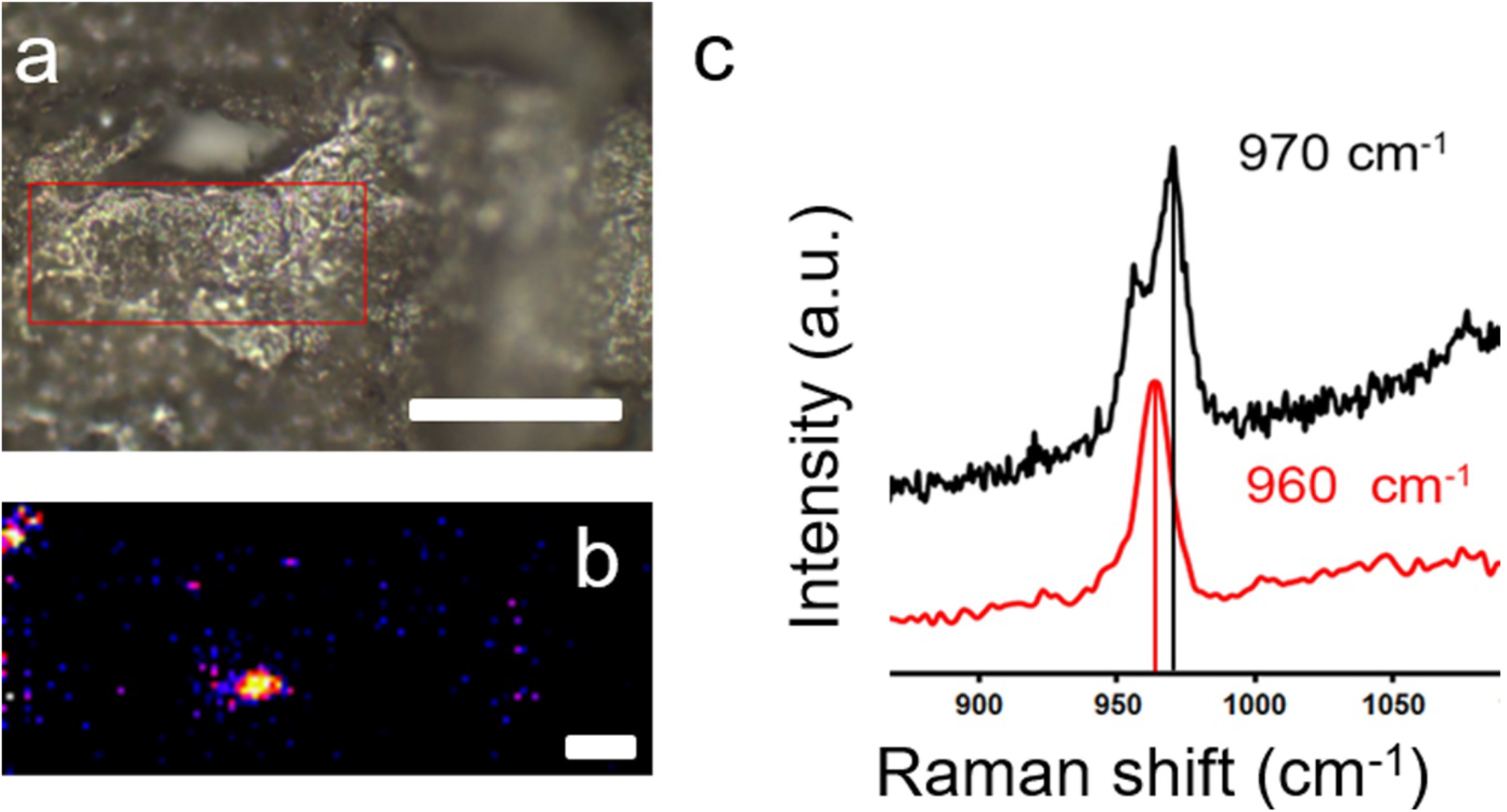
Optical micrograph and Raman mapping of 970 cm^-1^ peak on blood vessel wall. **(a)** Optical micrograph of a vessel wall, and region mapped with Raman indicated in red. Scale bar = 300 µm. **(b)** Raman 970 +/- 20 cm^-1^ peak map suggesting the presence of magnesium whitlockite spherical particles in the tissue. Scale bar = 4 µm. **(c)** Raman spectrum obtained from vessel wall for spherical particles (black spectrum) and large calcification (red spectrum).

The presence of hydroxyapatite^9,24,25^ and whitlockite^25-27^ has been acknowledged previously in breast tissues; however, they have never been identified as separate entities. Moreover, to the best of our knowledge, this is the first time calcification in breast tissue is shown to have the clear morphology of spherical particles of an average diameter of 434 ± 228 nm.

The difference in scale between the two types of minerals prompt us to suggest that the large minerals are likely the calcification observed on clinical mammograms and histopathological evaluation. In contrast, the small size of the spherical particles might mean that these are going undetected during clinical assessment.

The potential relationship between invasive breast malignancies and the spherical particles was further evaluated with an analysis of the characteristics of the particles. Initially, no association between particle size, type of tumour or grade was found (Fig. 8a). However, particle size in grade 1 tumours was found to be slightly larger than in grade 2 and grade 3 tumours (Fig. 8b).

**Figure 8:**
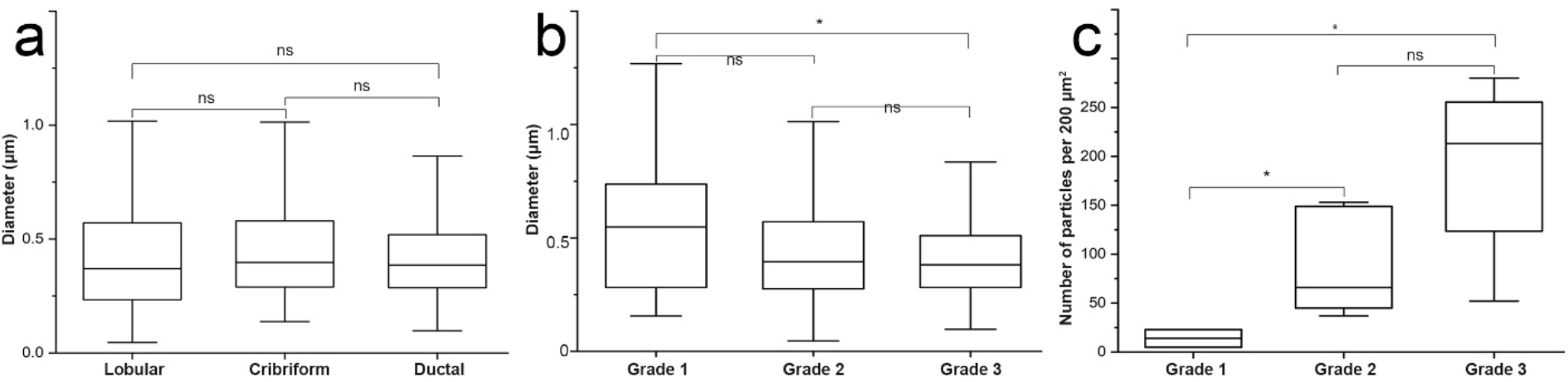
Dimensions and number of spherical particles on vasculature of invasive malignant breast tumours. (**a**) Particle diameter for different subtypes of invasive malignancy (lobular, cribriform and ductal (n = 300)) (mean ± SD). (**b**) Particle diameter in samples with different grades of malignancy (grade 1, grade 2 and grade 3 (n = 30, n = 600, n = 600)) (mean ± SD). *p =0.0294. (**c**) Spatial density distribution of particles in relation to different grades of malignancy (grade 1, grade 2 and grade 3 (n = 2, n = 5, n = 5)) (mean ± SD). One-way ANOVA with Dunnett’s T3 multiple comparisons post-hoc test. *p = 0.0298.

Conversely, the number of particles found in grade 1 tumours was significantly lower than in grade 2 and grade 3 tumours (Fig. 8c), indicating that these particles may accumulate over time, as the tumour progresses.

The association found between spherical particles and invasive malignant breast tumours could indicate a possible role in tumour development and metastasis. When found in soft tissues, calcium phosphate may have a significant impact on the recruitment and transdifferentiation of local or circulating cells^28-30^ through mechanobiological signalling, as stiffer matrices have been shown to have remarkable effects on cells^31^. It is well documented that calcium phosphates may induce apoptosis in human breast cancer cells^32^, while hydroxyapatite of different carbonate contents may considerably affect breast cancer cell behavior^33^. Moreover, the size of calcium phosphate crystals was also found to increase cell adhesion, whereas an increase in mineral crystallinity was reported to have the opposite effect^34^. Also, an increase in crystallinity has been correlated to the increase of osteolytic protein secretion from breast cancer cells^34,35^, which has in turn been correlated to bone metastasis^35^.

In spite of all the biological effects that may be caused by the presence of spherical particles in invasive malignant breast tumours, their origins are still elusive. Recent work in the field of cardiovascular^36^ and ocular^37^ mineralization has reported the presence of similar particles, whose origins are still debated^37-39^.

Our finding of spherical particles in invasive malignant breast tumours also suggests there may be a new calcification mechanism, the study of which could potentially contribute to a better understanding of tumour biology. Future studies in the field might benefit from exploring the strong association between these spherical particles and invasive malignant breast tumours. An investigation of the effect of these particles on breast cancer cells could enhance our understanding of their effect on tumour microenvironment, progression and metastasis. Finally, including their specific physicochemical properties in such studies, as has been done for the hydroxyapatite mineral, would add to our knowledge about the roles played by different materials in breast cancer.

## Acknowledgements

The authors would like to thank Shweta Agarwal for her assistance with FIB and TEM analyses. We also thank Charalambos Kallepitis for early discussions and suggestions for particle size measurement. We gratefully acknowledge support from the Rosetrees Trust (A1153). The authors wish to acknowledge the roles of the Breast Cancer Now Tissue Bank in collecting and making available samples and/or data, and of the patients who generously donated their tissues and shared their data to be used in the generation of this publication. We would also like to thank Francesca Launchbury and Angela Richard-Londt, from the UCL IQPath Institute of Neurology, for their help with histological analyses; Alba Rodríguez Meira, for her help with sample preparation.

## Author contributions

E.T. and W.D. performed sample preparation. E.T., W.D., and S.B. conducted microscopy work. J. O. and I.K.H. conducted Raman spectroscopy. E.T. evaluated the spatial distribution of particles. E.T., A.O., L.M., and S.B. designed the study. E.T., A.O., I.K.H., L.M., and SB contributed to scientific discussions, data interpretation and to the writing of the manuscript.

