## Supplementary info for "Nano-analytical Characterization of Breast Tissue Biopsies Reveals an Association between Spherical Nano and Micro Particles and Invasive Malignant Breast Tumours"

**Table S1:** Diagnostic categorisation of breast tissue samples.

| **Type of tissue** | **Number of cases** |
|---|---|
| Healthy tissue | 11 |
| Fibroadenoma | 14 |
| Benign Phyllodes tumour | 1 |
| Fibrotic change and cysts | 1 |
| Fibromatosis | 1 |
| Fibroepithelial lesion | 1 |
| Benign complex sclerosing papillary lesion | 1 |
| Hamartoma | 1 |
| Sclerosing adenosis | 1 |
| Uncapsulated mass with slightly disorganised glands and pseudo angiomatous stromal hyperplasia favouring Hamartoma | 1 |
| Paget's disease | 1 |
| Borderline Phyllodes | 1 |
| Ductal Carcinoma in Situ | 2 |
| Invasive Ductal Carcinoma | 16 |
| Invasive Cribriform Carcinoma | 7 |
| Invasive Lobular Carcinoma | 8 |
| Invasive carcinoma of unknown type | 14 |

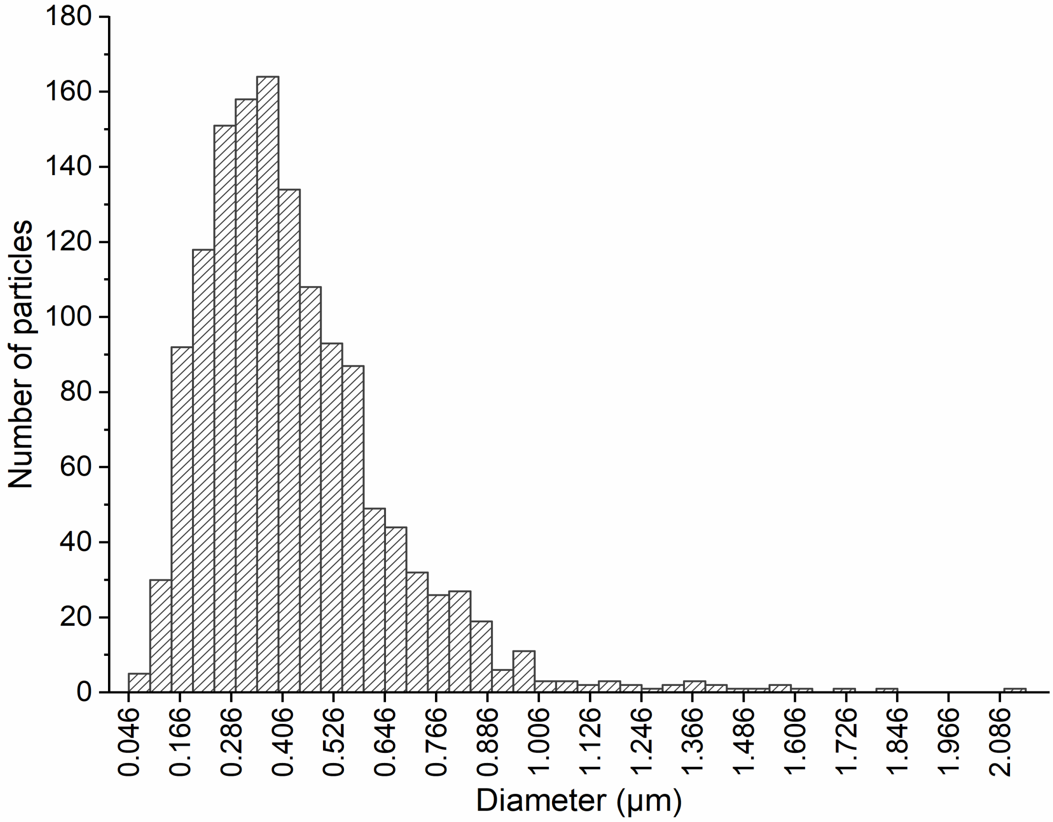


**Figure S1:** Size distribution of 1400 spherical particles from invasive malignant breast tumour biopsies. The smallest particle had a diameter of 46 nm and the largest a diameter of 2.15 µm, with the average size being 434 nm and a standard deviation of ± 228 nm.

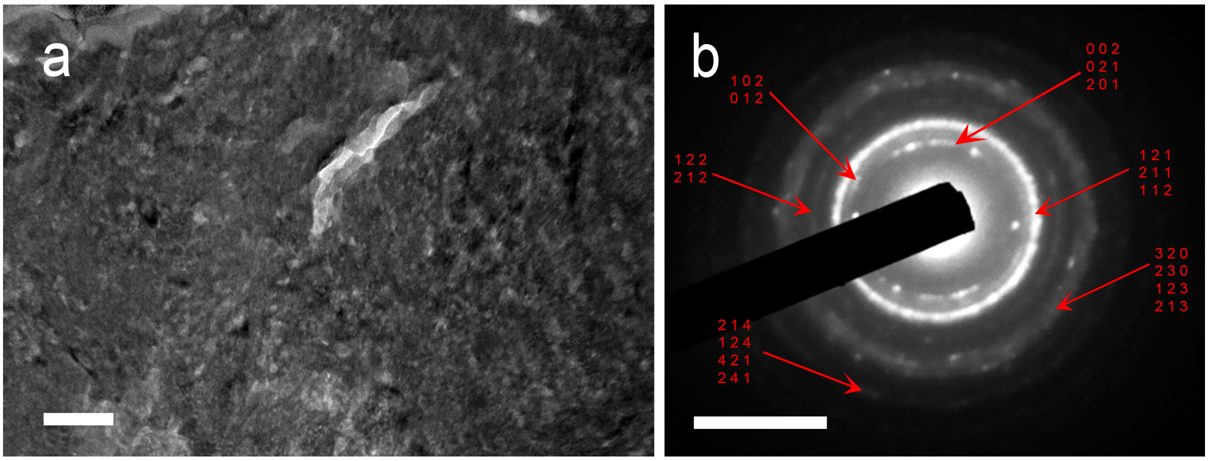


**Figure S2:** TEM micrograph (scale bar = 100 nm) and SAED (scale bar = 5 $nm^{-1}$) of large mineral observed in a benign phyllodes tumour.

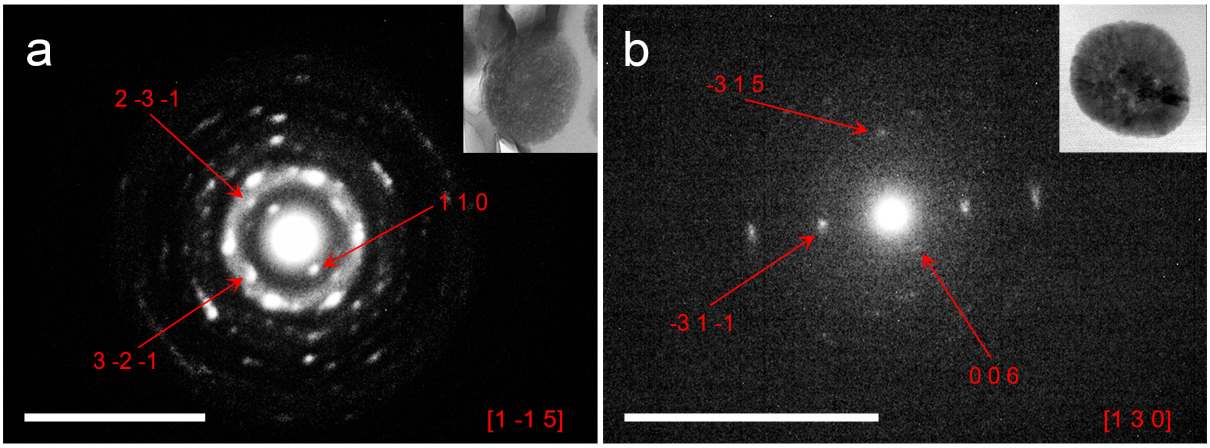


**Figure S3: TEM-SAED micrograph of a FIB-prepared section of a spherical particle.** (**a**) Representative electron diffraction pattern obtained from a spherical particle, showing the vector points (1 1 0), (-3 1 5) and (-3 1 -1) corresponding to the zone axis [1 -1 5] of magnesium whitlockite. (**b**) Representative electron diffraction pattern obtained from a spherical particle, showing the vector points (0 0 6), (2 -3 -1) and (3 -2 -1), corresponding to the zone axis [1 3 0] of magnesium whitlockite. Scale bar = 10 $nm^{-1}$.

### Experimental procedures

*Sample preparation*

A total of eighty-one samples were analysed (sample information presented on Supplementary Table S1). Samples were obtained from the Department of Surgery and Cancer at the Imperial Centre for Translational and Experimental Medicine (UK), Breast Cancer Now Tissue Bank (UK), and the Cantonal Hospital of St. Gallen (Switzerland). For sample collection, informed signed consent was obtained from either the patient or their next of kin. Samples were processed, analysed and disposed of according to the ethical approval obtained.

*Histology*

Histopathological slides of breast cancer biopsies were produced using standard preparation procedures. The tissue was formalin-fixed, paraffin-embedded and cut into sections of 3-5 μm in thickness, using a microtome. Haematoxylin and Eosin (H&E) staining was carried out on the slides, which were scanned using a Leica SCN400 scanner and a VS120 Olympus microscope.

*Scanning Electron Microscopy*

For scanning electron microscopy (SEM) analysis, consecutive histological slides were used. Dewaxing with pure xylene was carried out to enable tissue analysis, as follows. The samples were immersed for three times in pure xylene solutions at 5-minute intervals. Samples were then painted with silver conductive paint and coated with a 10 nm carbon layer using a Quorum K975X coater, prior to imaging. A Hitachi S-3499N and a Carl Zeiss Crossbeam microscopes were used for imaging, with both the secondary electron (SE) and backscattering electron (BSE) modes. Using both imaging modalities, density-dependent images (DDC-SEM) were also acquired, as previously described [24]. Energy-dispersive X-ray spectroscopy (EDX) analysis was carried out using Oxford Instruments EDX detectors, integrated into both microscopes. For the analysis, accelerating voltages of 5 kV and 10 kV and a working distance of 10 mm were used.

*Focused Ion Beam*

FEI Helios NanoLab 600 Dual Beam Focused Ion Beam (FIB) System was used in preparation for TEM imaging. A micro-region of each sample was coated with a Platinum layer at 30 kV and 93 pA. Following that, currents between 93 pA and 2.8 nA were used for section lift out and thinning to 100 nm.

*Transmission Electron Microscopy*

FIB prepared sections were imaged using a JEOL JEM 2100Plus Transmission Electron Microscope at 200 kV and 120 kV.

EDX analysis and selected area electron diffraction (SAED) were also carried out. Indexing of the patterns was carried out manually, using the measurement tools on Gatan Microscopy Suite 2 software. All data collected was then compared to existing available data on the CrystalWorks Server (https://cds.dl.ac.uk/).

*Raman spectroscopy*

Raman spectral imaging was performed on a WITec alpha300R Confocal Raman Microscope (WITec, Ulm, Germany) using de-paraffinised histology sections that had been previously SEM imaged, in order for areas of interest to be identified. A diode laser, 532 nm, 75 mW, was used for excitation. The laser beam was focused through a 50x objective (NA 0.8, Zeiss EC "Epiplan-Neofuar" DIC, Zeiss, Oberkochen, Germany). Raman signals were dispersed by grating (600 $gmm^{-1}$ for the mapping, and the single spectrum of the compact calcification (10 mW, 5s)), 1800 $gmm^{-1}$ for the spectra of the spherical particles (3 mW, 90 s) (spectrograph UHTS 300 for VIS, WITec, Ulm, Germany) and the spectra were acquired with a thermoelectrically cooled CCD detector. Control FOUR software (WITec) was used for measurement and Project FOUR Plus (WITec) for spectral data processing.

*Image analysis*

FIJI ImageJ software was used for all image analysis work. Images of suitable samples were used, taken at the same magnification and microscope settings. All physicochemical characterisation measurements (occurrence, size, spatial density) were carried out manually.

*Statistical analysis*

All statistical analyses were done using OriginLab 2019 and GraphPad Prism 8.3.1 software. A one-way Brown-Forsythe and Welch ANOVA with Dunnett's T3 multiple comparisons post hoc test (409) was used ($p<0.05$). All box blots represent the upper and lower quartiles; the whiskers indicate standard deviation, and the middle line the median value. All data is represented as the average value ± standard deviation.
